# Distance-to-optimum biological drift as a new framework for interpreting routine laboratory results: a benchmark against Reference Change Values across 62 routine biomarkers

**DOI:** 10.64898/2026.07.06.736744

**Authors:** Clément Bézier, Jakez Rolland, Ronan Boutin, Damien Gruson

## Abstract

**Background:** We propose the biological drift framework for the interpretation of biological test results: a z-score-like framework based on optimized and personalized reference populations and a distance-to-optimum drift metric for longitudinal interpretation relative to an estimated individual optimum. We benchmarked biological drifts against Reference Change Values (RCVs), which are used to interpret serial laboratory results by defining the minimum change expected to exceed normal within-subject biological variation (*CVi*).

**Objectives:** To benchmark biological drifts against the classical biological-variation framework and assess their consistency with RCV thresholds across routine biomarkers.

**Methods:** For 62 routine biomarkers, biological drift levels were compared with RCVs after transformation to test the consistency between the two frameworks.

**Results:** Severe biological drifts mostly exceeded the 95% RCV threshold, indicating changes unlikely to be explained by short-term biological variation alone. In contrast, moderate drifts reached the 95% RCV threshold for approximately one in two biomarkers, suggesting that many moderate distance-to-optimum deviations may remain within expected variability, particularly for biomarkers with large within-subject variation (*CVi*). Results are particularly interesting for the follow-up of people with diabetes and for the management of thyroid and hepatic disorders.

**Conclusions:** Biological drifts derived from optimized personalized reference populations are broadly consistent with the RCV framework for identifying biologically meaningful deviations from the optimum and may therefore be relevant for the monitoring of certain biomarkers across several medical conditions in clinical practice.

## Introduction

Personalized medicine requires clinicians and laboratory specialists to determine new thresholds based on sex, age, ethnicity, and other relevant criteria. This creates substantial complexity in routine patient management (1). In laboratory medicine, z-scores (standard deviation scores) provide a standardized way to interpret quantitative biomarkers relative to a reference population. By expressing results as deviations from a population mean, they enable harmonized interpretation across heterogeneous individuals and clinical contexts (1). Two main rationales support their widespread use:

(i) Addressing biological heterogeneity and enabling longitudinal assessment. Many biomarkers vary substantially with age, sex, and developmental stage, limiting the relevance of fixed thresholds. Z-scores address this issue by normalizing measurements against stratified reference populations, allowing both cross-sectional comparison and longitudinal monitoring while accounting for physiological changes. For insulin-like growth factor 1 (IGF-1), whose levels vary with age, z-scores enable discrimination between pathological and physiological variation and facilitate treatment monitoring (2, 3). Similarly, bone mineral density is interpreted using age- and sex-adjusted z-scores in younger populations to avoid misclassification and allow meaningful follow-up (4). Comparable approaches are used in childhood overweight and obesity, where z-scores account for growth-related variability (5), in age- and ethnicity-adjusted models to assess lung function via spirometry (6, 7), and for measurements of cardiac structures in infants, children, and adolescents (8). In this context, different z-score classes are sometimes used to grade severity.

(ii) Defining abnormality using statistical properties of reference populations. Z-scores also provide a direct statistical framework for defining abnormality. Under assumptions of normality, thresholds such as ± 1.96 standard deviations may correspond to the central 95% of the reference population, enabling consistent identification of outliers. This principle underlies their use in pediatric densitometry (9), in anthropometry to evaluate malnutrition in childhood (10), and in non-invasive assessment of trisomy 21 by DNA sequencing (11).

Alternatively, there is growing interest in person-centered interpretation, motivated by the observation that many measurands have an individual homeostatic set point and within-subject biological variation (*CV*_*i*_) that can be substantially smaller than between-subject variation (*CV*_*g*_) (12–14). Among these tools, the Reference Change Value (RCV) is routinely used to define the minimum significant difference between two consecutive results from the same individual, based on *CV*_*i*_ and analytical imprecision (*CV*_*a*_) (15, 16). Dedicated resources, including the European Federation of Clinical Chemistry and Laboratory Medicine (EFLM) biological variation database and standardized critical appraisal frameworks such as the Biological Variation Data Critical Appraisal Checklist (BIVAC), aim to increase transparency, quality, and comparability of biological variation studies (12, 13, 17).

In this context, we propose a z-score-inspired metric based on personalized reference values derived from healthy individuals sharing similar characteristics, such as age and sex. In previous work, we designed a new method to define optimized and personalized reference populations based on biological exclusion criteria in order to refine and personalize existing reference intervals (RIs). We showed that the resulting thresholds were more refined than the classical RIs analyzed (18). Based on these previous reference populations, we quantify “drifts” as distances from the estimated optimum (the corrected mean of the new reference population), using selected quantiles of the reference population distribution to obtain an explicit categorical stratification: a distance-to-optimum interpretation rather than a binary in-/out-of-range classification. This approach combines key aspects of both the z-score and the t-score (19): it preserves personalization by comparing individuals within homogeneous demographic groups, while using only healthy subjects as the reference population. As a result, each individual is benchmarked against the healthiest members of their own personalized category. We propose that in-range biological drifts have biological significance and that their interpretation may improve patient monitoring and care within conventional reference intervals.

Thus, standard z-scores (indicators of position relative to a reference), RCVs (indicators of short-term intra-patient variation), and distance-to-optimum drift (standardization against a reference population and longitudinal follow-up) form a triad for interpreting laboratory results.

The aim of the present study is to introduce the biological drift method and to benchmark these biological drifts against the classical biological-variation framework (RCV), in order to determine whether the magnitudes of deviations captured by biological drift are consistent with what would be considered a biologically significant change under the RCV framework. A key conceptual difference must be acknowledged: the RCV is defined for changes between two serial measurements (typically the most recent value compared with a previous value), whereas drift metrics quantify deviations relative to an estimated optimum value. Despite this difference in anchoring, we hypothesize that overall consistency should exist between (i) distance-to-optimum drifts (across predefined quantile levels) and (ii) exceedance of expected change thresholds (RCV) derived from *CV*_*i*_ and *CV*_*a*_. We therefore (1) formalized the distance-to-optimum drift metrics at several drift levels, (2) compared these drifts with RCVs computed at commonly used confidence levels, and (3) analyzed conditions under which the two frameworks converge or diverge. We analyzed 62 commonly measured biomarkers, and the objective was to evaluate the complementarity between the two approaches.

## Materials and Methods

### Study design and overview

This work is a retrospective methodological benchmarking study designed to compare (i) a classical biological-variation framework based on the Reference Change Value (RCV) and (ii) a set of “distance-to-optimum” drift metrics derived from optimized and personalized reference populations (18). For each biomarker, we quantified how far predefined drift thresholds were from the estimated individual optimum and assessed whether these deviations were consistent with changes that would be considered significant under the RCV framework at several confidence levels (80%, 90%, and 95%). Analyses were performed across 62 routinely measured biomarkers spanning clinical chemistry, endocrinology, hematology, and immunology panels.

### Data sources

Two data sources were combined:

1. **Biological variation estimates**. Median estimates of within-subject biological variation (*CV*_*i*_) (and, when available, between-subject variation) from the European Federation of Clinical Chemistry and Laboratory Medicine (EFLM) biological variation database (12).
2. **Optimized and personalized reference population outputs**. The corrected mean, obtained after Box-Cox transformation of the reference values, was considered the “personalized optimum”, whereas the quantiles {2.5%, 15.8%, 84.1%, 97.5%} and {68.4%, 95%} were used to stratify values for bilateral parameters (with both low and high thresholds) and unilateral parameters (with a high threshold only), respectively. For a given patient, clinicians can compare biological values against the deviation levels of a personalized reference population with the same subject-specific characteristics to determine their classification (see Table 1). Because a specific reference population is assigned to each personalized patient profile, we studied the different thresholds for each biomarker and each patient profile (sex, age, and ABO group).

**Table 1.**
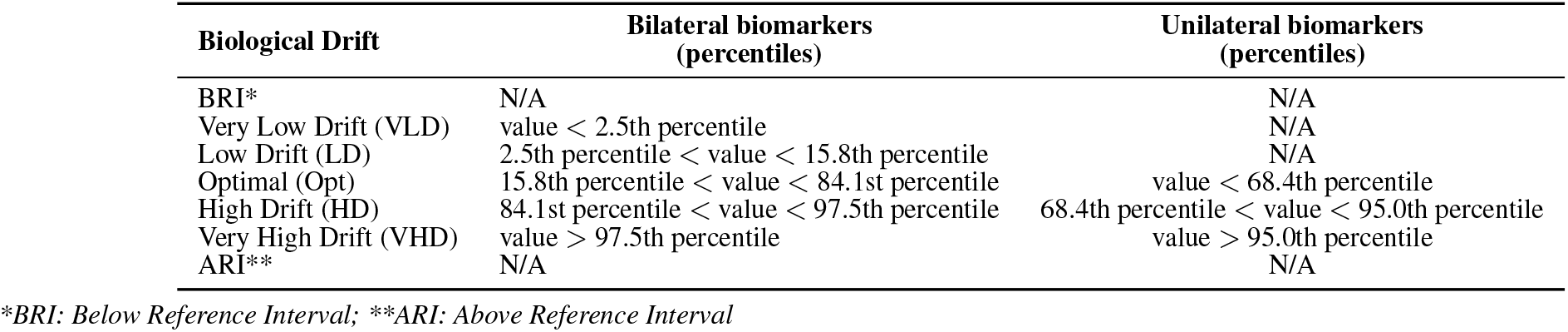
Classification of patients according to biological drift.

| Biological Drift | Bilateral biomarkers<br>(percentiles) | Unilateral biomarkers<br>(percentiles) |
| --- | --- | --- |
| BRI* | N/A | N/A |
| Very Low Drift (VLD) | value < 2.5th percentile | N/A |
| Low Drift (LD) | 2.5th percentile < value < 15.8th percentile | N/A |
| Optimal (Opt) | 15.8th percentile < value < 84.1st percentile | value < 68.4th percentile |
| High Drift (HD) | 84.1st percentile < value < 97.5th percentile | 68.4th percentile < value < 95.0th percentile |
| Very High Drift (VHD) | value > 97.5th percentile | value > 95.0th percentile |
| ARI** | N/A | N/A |

\*BRI: Below Reference Interval; \*\*ARI: Above Reference Interval

### Reference Change Value (RCV)

Because analytical variation (*CV*_*a*_) was not available for each measurand, we computed two-sided RCV thresholds at 95%, 90%, and 80% confidence as follows:

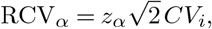

where *z*_*α*_ is the standard normal quantile corresponding to the desired two-sided confidence level (*z* = 1.96 for 95%, *z* = 1.645 for 90%, and *z* = 1.28 for 80%). RCVs were expressed as percentages. Biomarkers lacking *CV*_*i*_ (and therefore RCVs) were excluded from all RCV-versus-drift comparisons.

### Biological drift metrics

For each biomarker, we derived normalized deviations between the estimated optimum and four drift bounds corresponding to increasing levels of deviation:

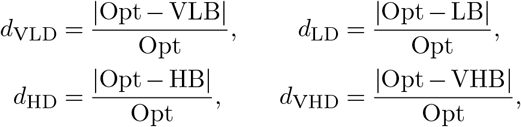

where Opt denotes the optimum estimate (mean of the reference population after Box-Cox normalization) and VLB/LB/HB/VHB denote the very-low, low, high, and very-high drift bounds, respectively. These quantities were computed as absolute relative differences (dimensionless) when divided by the optimum estimate (Opt) and subsequently converted to percentages.

We quantified the width of the “optimum” interval as a relative width:

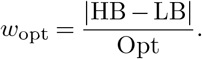

For biomarkers with only an upper bound and deviations toward higher values (C-reactive protein, D-dimer, alpha-fetoprotein, absolute basophil count, and carcinoembryonic antigen), we used a lower limit of 0, i.e.,

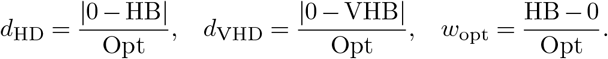

### Linking drift magnitudes to an “RCV-equivalent” confidence

Beyond comparing drift magnitudes with fixed RCV thresholds, we aimed to express each observed drift magnitude as an *RCV-equivalent confidence level*, i.e., the percentage of individuals for whom a deviation of that magnitude would still be considered compatible with normal within-subject biological variability (as summarized by *CV*_*i*_ under a Gaussian assumption). In the classical framework, RCV_95_ can be interpreted as the change magnitude that would be exceeded by only 5% of stable individuals when two consecutive measurements are compared; equivalently, 95% of stable individuals are expected to show absolute changes *below* this threshold, given *CV*_*i*_ (and the analytical component, if included). Here, we invert this logic for drift metrics: for a given deviation *d* (e.g., *d*_LD_, *d*_HD_, *d*_VHD_, or *w*_opt_), we compute the confidence level *p*_RCV_ such that RCV_*p*__RCV_ would match the observed drift magnitude. This provides a direct comparison on the same scale as standard RCV reporting: a large drift yields a high *p*_RCV_ (close to 1), meaning that only a small fraction of stable individuals would be expected to display such a deviation under the *CV*_*i*_ model, whereas a small drift yields a lower *p*_RCV_.

Formally, let *d* denote a deviation expressed as a fraction of the Opt value (e.g., *d*_LD_) and let *CV*_*i*_ be expressed as a per centage; the corresponding standardized statistic is:

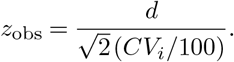

Let *X* ~ *N* (0, 1) represent patient variation standardized by the Opt value; the probability *P* (*X* ≤ *z*_obs_) represents the probability that a standard-normal fluctuation is smaller than the standardized observed deviation *z*_obs_. Therefore, the corresponding two-sided confidence (bounded to [0, 1]) can be written as:

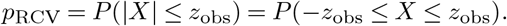

We refer to *p*_RCV_ as the “RCV percentage attained” by the deviation from the optimum; it can be interpreted as the fraction of stable individuals expected (under the Gaussian *CV*_*i*_ model) to have an absolute standardized change from the optimum that is smaller than the observed deviation magnitude.

## Results

### Global statistics across all biomarkers

For Opt–VHD and for Opt-width, no biomarker had a mean *p*_RCV_ below 95%. In contrast, mean *p*_RCV_ fell below 95% for Opt–VLD in only five biomarkers: serum iron, triglycerides, fasting insulin, absolute neutrophil count, and fibrinogen.

For the less severe (and therefore more frequent) Opt–LD and Opt–HD drifts, a larger number of biomarkers exhibited *p*_RCV_ *<* 95%. For Opt–LD, 33/57 biomarkers (58%) had *p*_RCV_ *<* 95%, 25/57 (44%) had *p*_RCV_ *<* 90%, and 11/57 (19%) had *p*_RCV_ *<* 80%. For Opt–HD, 28/62 biomarkers (45%) had *p*_RCV_ *<* 95%, 18/62 (29%) had *p*_RCV_ *<* 90%, and 3/62 (5%) had *p*_RCV_ *<* 80% (Figure 1).

**Fig. 1.**
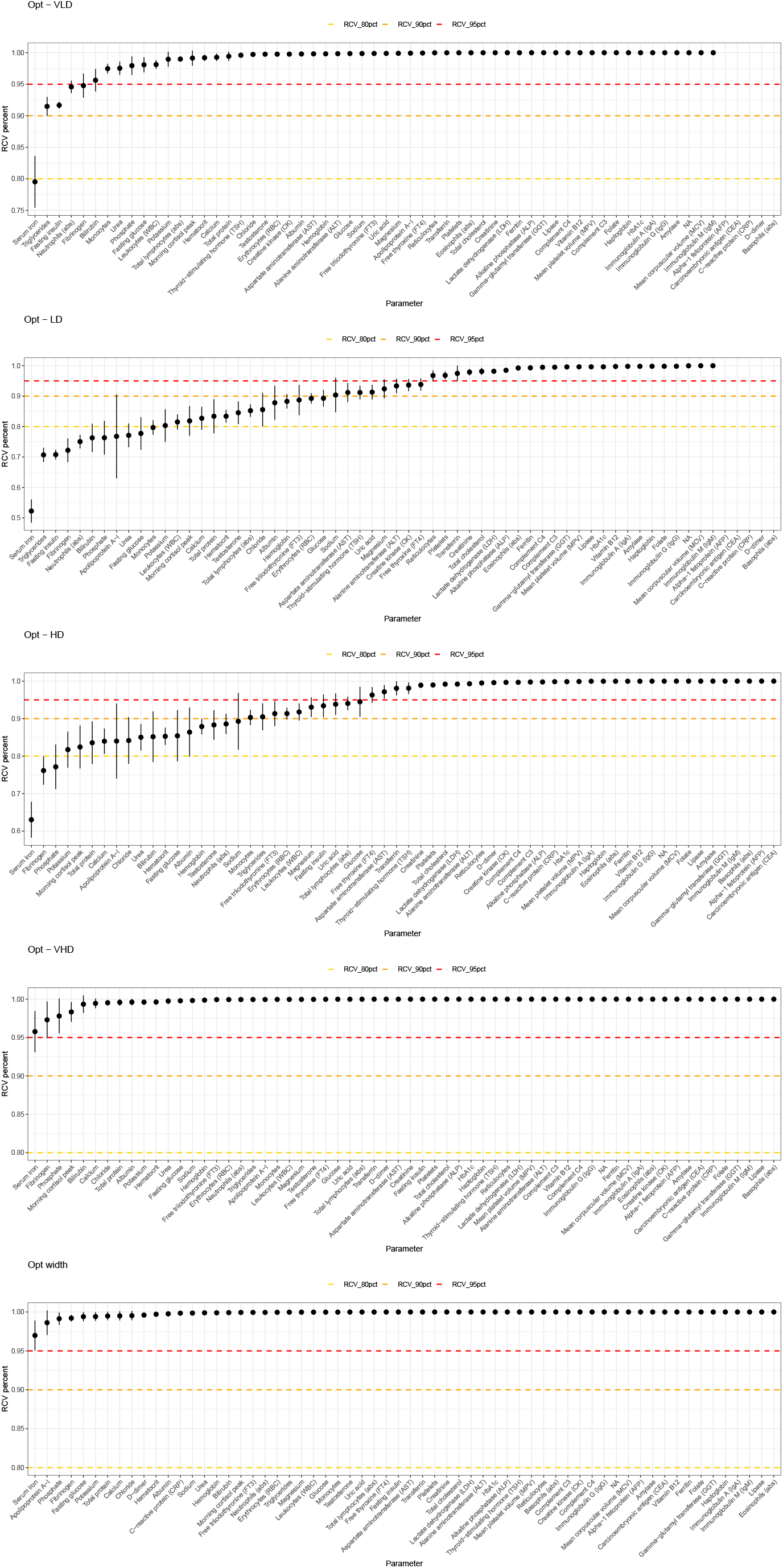
Comparison of *p*_RCV_ values (reported as RCV percent) for biological drift levels with the 80%, 90%, and 95% RCV thresholds. *p*_RCV_ values are expressed as means and standard deviations across all reference populations for a given biomarker.

To summarize these results across reference populations, we computed, for each biomarker and each drift level, the proportion of personalized reference-population models with *p*_RCV_ *>* 95%, and then averaged this proportion across biomarkers. On average, 91.76%, 44.67%, 53.37%, 98.82%, and 99.70% of reference-population models achieved *p*_RCV_ *>* 95% for Opt–VLD, Opt–LD, Opt–HD, Opt–VHD, and Opt-width, respectively.

The *p*_RCV_ values for each reference population included in the comparison were plotted for each biomarker in Supplementary Figures 1–5.

Finally, for each biomarker, we computed the 95th percentile of *p*_RCV_ across the set of personalized reference-population models (see Supplementary Table 1).

Biomarkers with the lowest *p*_RCV_ values (e.g., serum iron and bilirubin) appeared to have high *CV*_*i*_*/CV*_*g*_ ratios, meaning that *CV*_*i*_ is close to *CV*_*g*_ (see Figure 2).

**Fig. 2.**
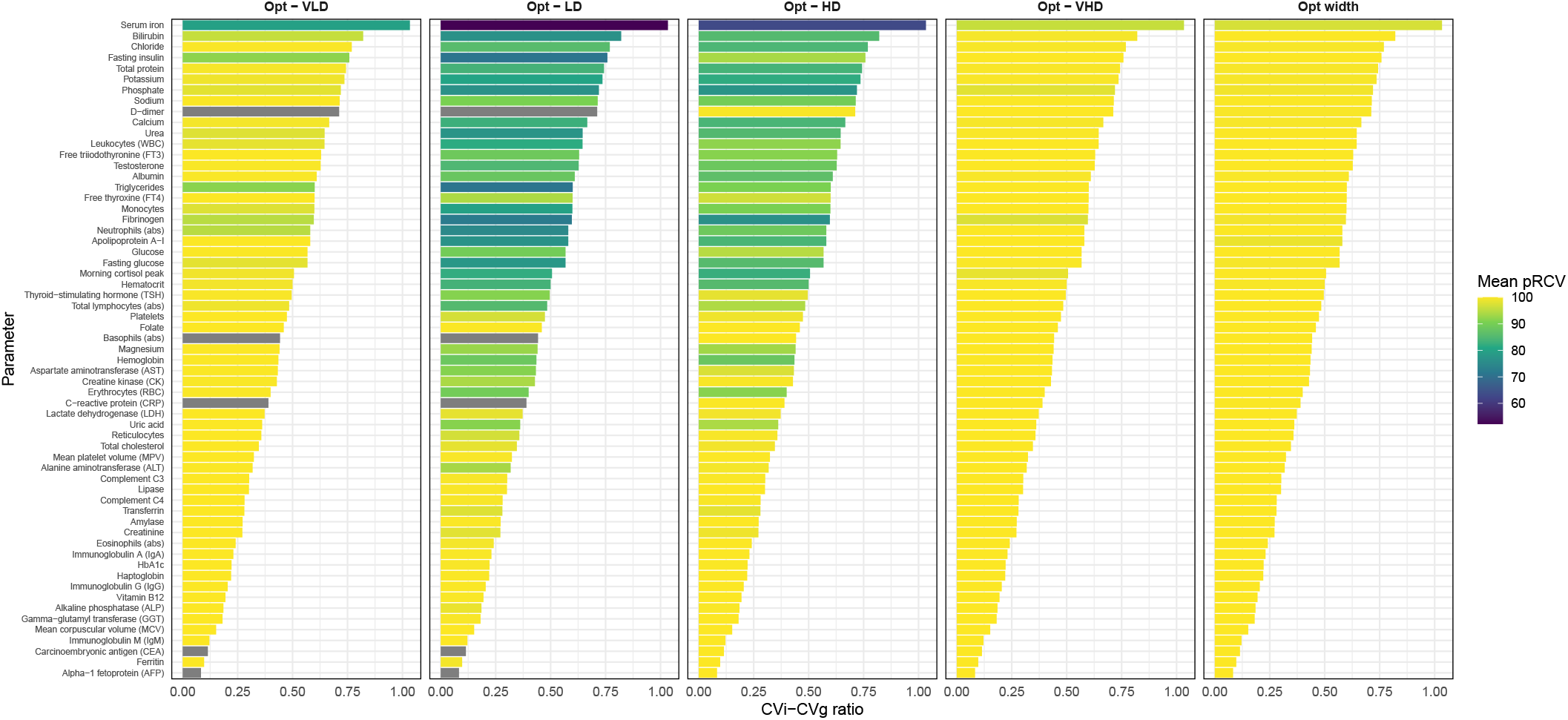
*CV*_*i*_*/CV*_*g*_ ratio and mean *p*_RCV_.

Overall, correlations were observed between the *p*_RCV_ values of drift deviations and the *CV*_*i*_*/CV*_*g*_ ratio (*p*-values from Pearson correlation tests *<* 10_*−*_4 for each drift type), with Pearson *r*^2^ values of 0.33, 0.73, 0.62, 0.25, and 0.31 for Opt– VLD, Opt–LD, Opt–HD, Opt–VHD, and Opt-width, respectively (see Figure 3).

**Fig. 3.**
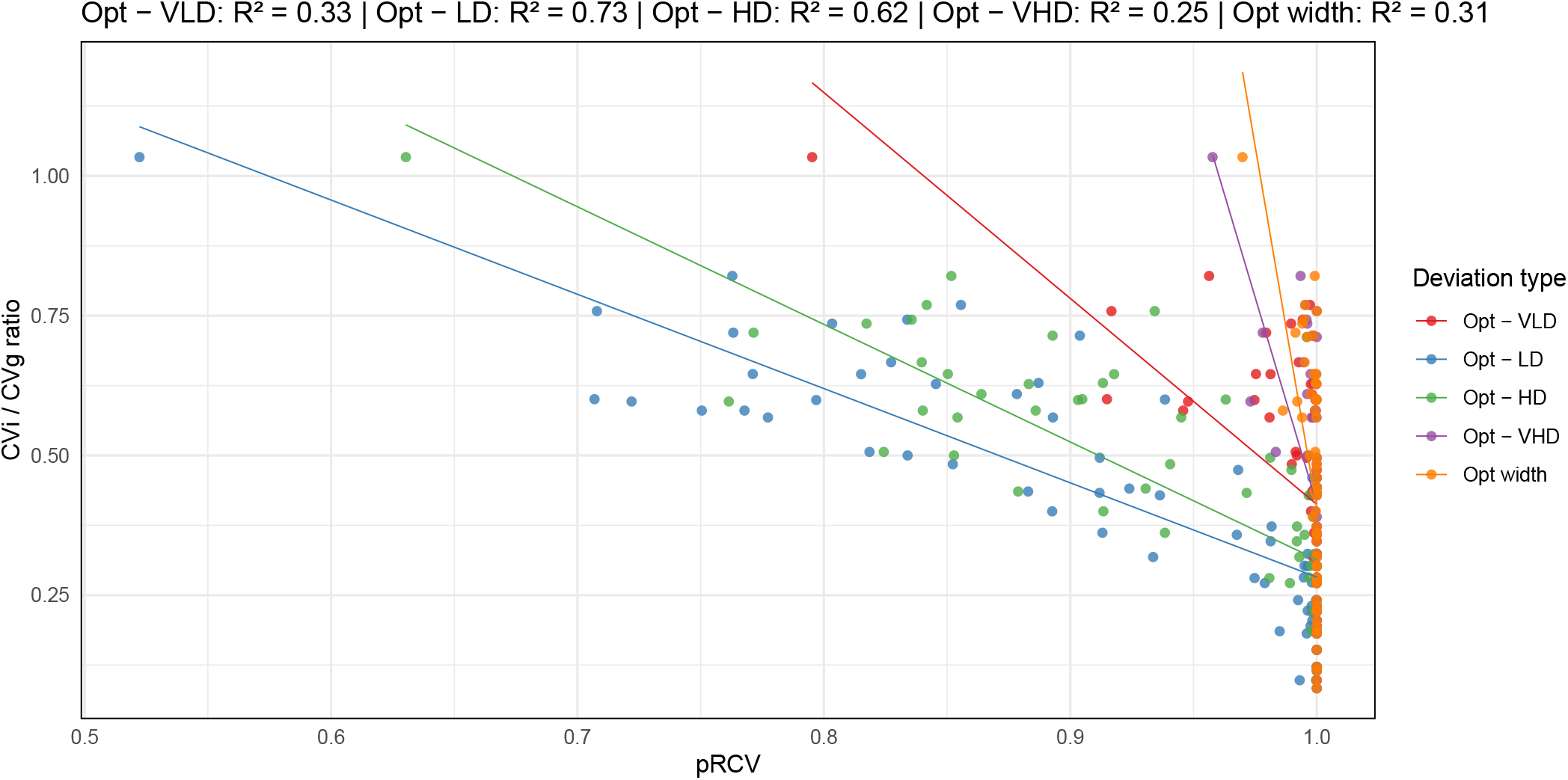
Correlation between *p*_RCV_ and the *CV*_*i*_*/CV*_*g*_ ratio.

### Illustrations for clinical situations of interest

For diabetes-related biomarkers, the 95th percentile of *p*_RCV_ was at least 86.87 for fasting glucose, 99.99 for HbA1c, 73.01 for fasting insulin, and 99.08 for creatinine (see Supplementary Table 1 and Supplementary Fig. 4). These findings suggest that LD/HD, VHD/VLD, and Opt-width deviations are generally consistent with clinically meaningful differences within the RCV framework. Overall, the LD–Opt distance tended to yield the lowest *p*_RCV_ values, suggesting an asymmetric distribution for several biomarkers.

For biomarkers used in the assessment of hypothyroidism and hyperthyroidism (free thyroxine, T4; free triiodothyronine, T3; and thyroid-stimulating hormone, TSH), LD–Opt and HD–Opt distances yielded *p*_RCV_ values close to 95%, whereas VHD–Opt, VLD–Opt, and Opt-width were associated with *p*_RCV_ values close to 100% (see Supplementary Table 1 and Supplementary Figs. 2 and 4).

For liver-related biomarkers, *p*_RCV_ values were close to 95% for AST and ALT in the LD category and exceeded 95% for the other drift classes. For lactate dehydrogenase (LDH), alkaline phosphatases (ALP), amylase, gamma-glutamyltransferase (GGT), and lipase, *p*_RCV_ values were consistently above 95% (see Supplementary Table 1 and Supplementary Figures 2 and 5).

## Discussion

These results highlight a marked asymmetry between extreme biological drifts (Very Low and Very High Drift) and moderate biological drifts (Low and High Drift) when interpreted through the lens of the Reference Change Value (RCV). For Opt–VHD (a deviation analogous to exceeding a reference interval) and for Opt-width, the mean *p*_RCV_ consistently remained above 95%, suggesting that these events are, for most measurands, larger than would be expected under the RCV framework for assessing the significance of serial changes (15, 16). In contrast, the frequent occurrence of Opt– LD and Opt–HD drifts with *p*_RCV_ *<* 95% for a substantial fraction of biomarkers indicates that many moderate deviations from the personalized optimum may fall within the “expected” short-term variability captured by current biological-variation estimates (12, 17, 20). This observation is consistent with the concept of individuality: when *CV*_*i*_ is relatively small, clinically meaningful changes can be detected early, whereas for biomarkers with larger *CV*_*i*_ values (close to *CV*_*g*_), moderate drifts may not exceed the RCV threshold and thus remain indistinguishable from noise (16, 21). This is particularly true for biomarkers that vary throughout the day or according to fasting status.

Importantly, the heterogeneity observed across personalized reference-population models, as reflected by the distribution and the 95th percentile of *p*_RCV_, suggests that within-population variability depends on personalization strata (e.g., age and sex). This heterogeneity can modify the probability that a given drift exceeds the RCV threshold, underscoring the need for a personalized perspective on biological variation (18).

Finally, because the performance of RCVs depends on the availability, quality, and applicability of the underlying *CV*_*i*_ and *CV*_*a*_ inputs (22), our findings reinforce the importance of using critically appraised biological-variation data (e.g., BIVAC-graded studies and curated repositories) when translating drift-based interpretation into clinical decision support (12, 13, 23). The new stratification thresholds derived from optimized personalized reference populations yielded results consistent with the RCV framework for the 62 biomarkers under study. When drift deviations fall below the 95% RCV threshold, this is frequently due to high *CV*_*i*_ values close to *CV*_*g*_, as demonstrated by the global correlation across all biomarkers.

Based on these results, distance-to-optimum drifts define value intervals within conventional reference intervals while remaining meaningful relative to the RCV framework.

A limitation of this study is that we did not have access to *CV*_*a*_, which can substantially affect RCV (22). This may have led to an underestimation of RCV and *p*_RCV_ when using *CV*_*i*_ alone.

From a clinical perspective, distance-to-optimum drifts in diabetes may therefore provide a useful complementary framework for diabetes diagnosis, glucose monitoring, assessment of comorbidities such as renal impairment, and evaluation of a return to the personalized optimum category after treatment or lifestyle modification, both in primary care and in hospital-based follow-up. For thyroid disorders, variations classified as VLD/LD, HD, or VHD are statistically significant and may be clinically informative. In some subclinical cases, TSH varies while free T4 and T3 remain within the conventional reference range. Distance-to-optimum drift relative to a personalized reference population adjusted for sex and age may therefore help detect such subtle abnormalities and refine the management of hypo- or hyperthyroidism. Treatment response could also be monitored more precisely through the return of TSH, free T4, and T3 toward the personalized optimum category. For liver assessment, distance-to-optimum drifts may be relevant for the personalized detection and longitudinal follow-up of non-alcoholic steatohepatitis (NASH). This approach may also have applications for prevention and for the identification of hepatic comorbidities associated with obesity or type 2 diabetes.

In conclusion, this study positions biological drift as a systematic tool for personalized medicine, as it standardizes all biomarker results against an optimized and personalized reference population free of biological abnormalities. It addresses the problem of multiple thresholds depending on factors such as sex and age, and goes further by taking sources of biological variation into account when defining the reference population. It therefore addresses both the need for a standardized interpretation model that takes into account patient’s natural progression (both clinical and biological) and the challenge of directly evaluating the statistical rarity of an individual patient’s result.

## Acknowledgements

We thank all contributors to the Bio Logbook project and all patients included in the different cohorts necessary for the design, development, and validation of the biological drift method.

## Contributions

CB carried out the analysis and wrote the article. JR and RB initiated this project and contributed to the analysis of the results and to the scientific strategy. JR contributed to the writing of the article. DG guided the discussion by aligning our methodologies with the current state of the art in clinical biology, analyzed the results, and contributed to the writing of the article.

## Conflict of interest

RB is the founder of Bio Logbook, and this did not affect the design, data collection, analysis, conclusions, decision to publish, or preparation of the manuscript. All other authors declare no potential conflicts of interest.

## Funding

NA.

## Bibliography

1. Roland RJ van Kimmenade, Marlies Kempers, Menko-Jan de Boer, Bart L Loeys, and Janneke Timmermans. A clinical appraisal of different z-score equations for aortic root assessment in the diagnostic evaluation of marfan syndrome. Genetics in Medicine, 15(7): 528–532, 2013.

2. Martin Bidlingmaier, Nele Friedrich, Rebecca T Emeny, Joachim Spranger, Ole D Wolthers, Josefine Roswall, Antje Körner, Barbara Obermayer-Pietsch, Christoph Hübener, Jovanna Dahlgren, et al. Reference intervals for insulin-like growth factor-1 (igf-i) from birth to senescence: results from a multicenter study using a new automated chemiluminescence igf-i immunoassay conforming to recent international recommendations. The Journal of Clinical Endocrinology & Metabolism, 99(5):1712–1721, 2014.

3. David R Clemmons. Consensus statement on the standardization and evaluation of growth hormone and insulin-like growth factor assays. Clinical chemistry, 57(4):555–559, 2011.

4. Joao Lindolfo Cunha Borges, Marina Sousa da Silva, Robert J Ward, Kathryn M Diemer, Swan S Yeap, and E Michael Lewiecki. Repeating vertebral fracture assessment: 2019 iscd official position. Journal of Clinical Densitometry, 22(4):484–488, 2019.

5. Tim J Cole, Mary C Bellizzi, Katherine M Flegal, and William H Dietz. Establishing a standard definition for child overweight and obesity worldwide: international survey. Bmj, 320 (7244):1240, 2000.

6. Philip H Quanjer, Sanja Stanojevic, Tim J Cole, Xaver Baur, Graham L Hall, Bruce H Culver, Paul L Enright, John L Hankinson, Mary SM Ip, Jinping Zheng, et al. Multi-ethnic reference values for spirometry for the 3–95-yr age range: the global lung function 2012 equations, 2012.

7. Graham L Hall, Nicole Filipow, Gregg Ruppel, Tolu Okitika, Bruce Thompson, Jane Kirkby, Irene Steenbruggen, Brendan G Cooper, Sanja Stanojevic, Bert Arets, et al. Official ers technical standard: Global lung function initiative reference values for static lung volumes in individuals of european ancestry. European Respiratory Journal, 57(3), 2021.

8. Michael D Pettersen, Wei Du, Mary Ellen Skeens, and Richard A Humes. Regression equations for calculation of z scores of cardiac structures in a large cohort of healthy infants, children, and adolescents: an echocardiographic study. Journal of the American Society of Echocardiography, 21(8):922–934, 2008.

9. Berta Magallares, Jorge Malouf, Helena Codes-Méndez, Hye Sang Park, Jocelyn Betancourt, Gloria Fraga, Estefanía Quesada-Masachs, Mireia López-Corbeto, Montserrat Torrent, Ana Marín, et al. Pediatric densitometry: is the z score adjustment necessary in all cases? Frontiers in endocrinology, 16:1587382, 2025.

10. World Health Organization et al. Recommendations for data collection, analysis and reporting on anthropometric indicators in children under 5 years old. 2019.

11. Rossa WK Chiu, Ranjit Akolekar, Yama WL Zheng, Tak Y Leung, Hao Sun, KC Allen Chan, Fiona MF Lun, Attie TJI Go, Elizabeth T Lau, William WK To, et al. Non-invasive prenatal assessment of trisomy 21 by multiplexed maternal plasma dna sequencing: large scale validity study. Bmj, 342, 2011.

12. A. K. Aarsand, A. Carobene, S. Sandberg, et al. The eflm biological variation database. Clinical Chemistry and Laboratory Medicine, 2021.

13. A. K. Aarsand, S. Sandberg, et al. The eflm biological variation data critical appraisal checklist (bivac): a standardized tool to evaluate studies on biological variation. Clinical Chemistry and Laboratory Medicine, 2018.

14. Eirik Åsen Røys, Kristin Viste, Christopher-John Farrell, Ralf Kellmann, Bashir Alaour, Marit Sverresdotter Sylte, Janniche Torsvik, Heidi Strand, Michael Marber, Torbjørn Omland, et al. A parametric empirical bayes approach to personalized reference intervals and reference change values. Clinical Chemistry, 71(11):1147–1157, 2025.

15. C Ricos, F Cava, JV García-Lario, A Hernández, N Iglesias, CV Jimenez, J Minchinela, C Perich, M Simón, MV Domenech, et al. The reference change value: a proposal to interpret laboratory reports in serial testing based on biological variation. Scandinavian journal of clinical and laboratory investigation, 64(3):175–184, 2004.

16. C. G. Fraser. Biological Variation: From Principles to Practice. AACC Press, 2001.

17. Aasne K Aarsand, Jorge Díaz-Garzón, Pilar Fernandez-Calle, Elena Guerra, Massimo Locatelli, William A Bartlett, Sverre Sandberg, Thomas Røraas, Ferruccio Ceriotti, Una ørvim Sølvik, et al. The eubivas: within-and between-subject biological variation data for eletrolytes, lipids, urea, uric acid, total protein, total bilirubin, direct bilirubin, and glucose. Clinical chemistry, 64(9):1380–1393, 2018.

18. Ronan Boutin, Jakez Rolland, Marie Codet, Clément Bézier, Nathalie Maes, Philippe Kolh, Leila Equinet, Marie Thys, Michel Moutschen, Pierre-Jean Lamy, et al. Use of hospital big data to optimize and personalize laboratory test interpretation with an application. Clinica Chimica Acta, 561:119763, 2024.

19. Hans P Dimai. Use of dual-energy x-ray absorptiometry (dxa) for diagnosis and fracture risk assessment; who-criteria, t-and z-score, and reference databases. Bone, 104:39–43, 2017.

20. Gallum G Fraser and Eugene K Harris. Generation and application of data on biological variation in clinical chemistry. Critical reviews in clinical laboratory sciences, 27(5):409–437, 1989.

21. Per Hyltoft Petersen, Esther Jensen, Carmen Ricós, Per E Jørgensen, and Natàlia Iglesias Canadell. Reference change values and power functions. Clinical Chemistry & Laboratory Medicine, 42(4), 2004.

22. Callum G Fraser. Reference change values. Clinical Chemistry & Laboratory Medicine, 50 (5), 2012.

23. Sverre Sandberg, Callum G Fraser, Andrea Rita Horvath, Rob Jansen, Graham Jones, Wytze Oosterhuis, Per Hyltoft Petersen, Heinz Schimmel, Ken Sikaris, and Mauro Panteghini. Defining analytical performance specifications: consensus statement from the 1st strategic conference of the european federation of clinical chemistry and laboratory medicine. Clinical Chemistry and Laboratory Medicine (CCLM), 53(6):833–835, 2015.

