## Supplementary Material for "Distance-to-optimum biological drift as a new framework for interpreting routine laboratory results: a benchmark against Reference Change Values across 62 routine biomarkers"

**Supplementary Table. 1:** Quantile 95% of the distribution of $p_{\mathrm{RCV}}$ for each parameter and deviation level.

| Parameter | Deviation | Quantile 95% |
|---|---|---|
| Alanine aminotransferase (ALT) | Opt-HD | 99.9384 |
| Alanine aminotransferase (ALT) | Opt-LD | 97.0216 |
| Alanine aminotransferase (ALT) | Opt-VLD | 99.9721 |
| Alanine aminotransferase (ALT) | Opt Width | 100 |
| Albumin | Opt-HD | 95.0373 |
| Albumin | Opt-LD | 94.8815 |
| Albumin | Opt-VLD | 99.9988 |
| Albumin | Opt Width | 99.9941 |
| Alpha-1 fetoprotein (AFP) | Opt-HD | 100 |
| Alpha-1 fetoprotein (AFP) | Opt Width | 100 |
| Amylase | Opt-HD | 99.9998 |
| Amylase | Opt-LD | 99.8 |
| Amylase | Opt-VLD | 100 |
| Amylase | Opt Width | 100 |
| Carcinoembryonic antigen (CEA) | Opt-HD | 100 |
| Carcinoembryonic antigen (CEA) | Opt Width | 100 |
| Apolipoprotein A-I | Opt-HD | 96.5004 |
| Apolipoprotein A-I | Opt-LD | 97.0656 |
| Apolipoprotein A-I | Opt-VLD | 100 |
| Apolipoprotein A-I | Opt Width | 99.9991 |
| Aspartate aminotransferase (AST) | Opt-HD | 99.0362 |
| Aspartate aminotransferase (AST) | Opt-LD | 94.0878 |
| Aspartate aminotransferase (AST) | Opt-VLD | 99.9589 |
| Aspartate aminotransferase (AST) | Opt Width | 100 |
| Bilirubin | Opt-HD | 93.2566 |
| Bilirubin | Opt-LD | 81.3774 |
| Bilirubin | Opt-VLD | 97.0613 |
| Bilirubin | Opt Width | 100 |
| Calcium | Opt-HD | 88.1547 |
| Calcium | Opt-LD | 87.0734 |
| Calcium | Opt-VLD | 99.829 |
| Calcium | Opt Width | 99.8278 |
| Chloride | Opt-HD | 93.0022 |
| Chloride | Opt-LD | 92.5806 |
| Chloride | Opt-VLD | 99.9804 |
| Chloride | Opt Width | 99.9593 |
| Total cholesterol | Opt-HD | 99.7271 |
| Total cholesterol | Opt-LD | 99.5063 |
| Total cholesterol | Opt-VLD | 100 |
| Total cholesterol | Opt Width | 100 |
| Vitamin B12 | Opt-HD | 99.9998 |
| Vitamin B12 | Opt-LD | 99.8969 |
| Vitamin B12 | Opt-VLD | 100 |
| Vitamin B12 | Opt Width | 100 |
| Complement C3 | Opt-HD | 99.9082 |
| Complement C3 | Opt-LD | 99.8299 |
| Complement C3 | Opt-VLD | 100 |
| Complement C3 | Opt Width | 100 |
| Complement C4 | Opt-HD | 99.9593 |
| Complement C4 | Opt-LD | 99.7259 |
| Complement C4 | Opt-VLD | 100 |
| Complement C4 | Opt Width | 100 |
| Morning cortisol peak | Opt-HD | 87.7973 |
| Morning cortisol peak | Opt-LD | 87.3719 |
| Morning cortisol peak | Opt-VLD | 99.9466 |
| Morning cortisol peak | Opt Width | 99.9997 |
| Creatine kinase (CK) | Opt-HD | 99.9934 |
| Creatine kinase (CK) | Opt-LD | 96.6078 |
| Creatine kinase (CK) | Opt-VLD | 99.9464 |
| Creatine kinase (CK) | Opt Width | 100 |
| Creatinine | Opt-HD | 99.7873 |
| Creatinine | Opt-LD | 99.0809 |
| Creatinine | Opt-VLD | 100 |
| Creatinine | Opt Width | 100 |
| C-reactive protein (CRP) | Opt-HD | 99.9446 |
| C-reactive protein (CRP) | Opt Width | 99.9446 |
| D-dimer | Opt-HD | 99.9269 |
| D-dimer | Opt Width | 99.9269 |
| Erythrocytes (RBC) | Opt-HD | 93.8397 |
| Erythrocytes (RBC) | Opt-LD | 91.5235 |
| Erythrocytes (RBC) | Opt-VLD | 99.9075 |
| Erythrocytes (RBC) | Opt Width | 99.9889 |
| Ferritin | Opt-HD | 100 |
| Ferritin | Opt-LD | 99.9077 |
| Ferritin | Opt-VLD | 99.9997 |
| Ferritin | Opt Width | 100 |
| Serum iron | Opt-HD | 70.2462 |
| Serum iron | Opt-LD | 57.7465 |
| Serum iron | Opt-VLD | 85.9111 |
| Serum iron | Opt Width | 99.2725 |
| Fibrinogen | Opt-HD | 80.7088 |

| Parameter | Deviation | Quantile 95% |
|---|---|---|
| Fibrinogen | Opt-LD | 77.6845 |
| Fibrinogen | Opt-VLD | 97.9106 |
| Fibrinogen | Opt Width | 99.6563 |
| Folate | Opt-HD | 100 |
| Folate | Opt-LD | 99.9673 |
| Folate | Opt-VLD | 100 |
| Folate | Opt Width | 100 |
| Gamma-glutamyl transferase (GGT) | Opt-HD | 100 |
| Gamma-glutamyl transferase (GGT) | Opt-LD | 99.9486 |
| Gamma-glutamyl transferase (GGT) | Opt-VLD | 100 |
| Gamma-glutamyl transferase (GGT) | Opt Width | 100 |
| Fasting glucose | Opt-HD | 94.5123 |
| Fasting glucose | Opt-LD | 86.8709 |
| Fasting glucose | Opt-VLD | 99.6665 |
| Fasting glucose | Opt Width | 99.9859 |
| Glucose | Opt-HD | 98.1788 |
| Glucose | Opt-LD | 93.0576 |
| Glucose | Opt-VLD | 99.9819 |
| Glucose | Opt Width | 99.9996 |
| Haptoglobin | Opt-HD | 99.9644 |
| Haptoglobin | Opt-LD | 99.8986 |
| Haptoglobin | Opt-VLD | 100 |
| Haptoglobin | Opt Width | 100 |
| Hematocrit | Opt-HD | 88.376 |
| Hematocrit | Opt-LD | 86.6911 |
| Hematocrit | Opt-VLD | 99.6424 |
| Hematocrit | Opt Width | 99.8907 |
| HbA1c | Opt-HD | 100 |
| HbA1c | Opt-LD | 100 |
| HbA1c | Opt-VLD | 100 |
| HbA1c | Opt Width | 100 |
| HbA1c | Opt-HD | 99.9999 |
| HbA1c | Opt-LD | 99.9995 |
| HbA1c | Opt-VLD | 100 |
| HbA1c | Opt Width | 100 |
| Hemoglobin | Opt-HD | 90.8508 |
| Hemoglobin | Opt-LD | 90.6765 |
| Hemoglobin | Opt-VLD | 99.9865 |
| Hemoglobin | Opt Width | 99.9599 |
| Immunoglobulin A (IgA) | Opt-HD | 99.9949 |
| Immunoglobulin A (IgA) | Opt-LD | 99.83 |
| Immunoglobulin A (IgA) | Opt-VLD | 100 |
| Immunoglobulin A (IgA) | Opt Width | 100 |
| Immunoglobulin G (IgG) | Opt-HD | 99.9918 |
| Immunoglobulin G (IgG) | Opt-LD | 99.9441 |
| Immunoglobulin G (IgG) | Opt-VLD | 100 |
| Immunoglobulin G (IgG) | Opt Width | 100 |
| Immunoglobulin M (IgM) | Opt-HD | 100 |
| Immunoglobulin M (IgM) | Opt-LD | 100 |
| Immunoglobulin M (IgM) | Opt-VLD | 100 |
| Immunoglobulin M (IgM) | Opt Width | 100 |
| Fasting insulin | Opt-HD | 97.5393 |
| Fasting insulin | Opt-LD | 73.0102 |
| Fasting insulin | Opt-VLD | 92.356 |
| Fasting insulin | Opt Width | 100 |
| Lactate dehydrogenase (LDH) | Opt-HD | 99.8453 |
| Lactate dehydrogenase (LDH) | Opt-LD | 98.7963 |
| Lactate dehydrogenase (LDH) | Opt-VLD | 99.9993 |
| Lactate dehydrogenase (LDH) | Opt Width | 100 |
| Leukocytes (WBC) | Opt-HD | 94.6729 |
| Leukocytes (WBC) | Opt-LD | 85.7913 |
| Leukocytes (WBC) | Opt-VLD | 98.9173 |
| Leukocytes (WBC) | Opt Width | 99.9985 |
| Lipase | Opt-HD | 100 |
| Lipase | Opt-LD | 99.9096 |
| Lipase | Opt-VLD | 100 |
| Lipase | Opt Width | 100 |
| Total lymphocytes (abs) | Opt-HD | 96.1132 |
| Total lymphocytes (abs) | Opt-LD | 87.6017 |
| Total lymphocytes (abs) | Opt-VLD | 99.5314 |
| Total lymphocytes (abs) | Opt Width | 99.9997 |
| Magnesium | Opt-HD | 96.0749 |
| Magnesium | Opt-LD | 96.3361 |
| Magnesium | Opt-VLD | 100 |
| Magnesium | Opt Width | 99.9991 |
| Monocytes | Opt-HD | 93.7649 |
| Monocytes | Opt-LD | 82.6062 |
| Monocytes | Opt-VLD | 98.311 |
| Monocytes | Opt Width | 99.9976 |
| Alkaline phosphatase (ALP) | Opt-HD | 99.9813 |
| Alkaline phosphatase (ALP) | Opt-LD | 99.3552 |
| Alkaline phosphatase (ALP) | Opt-VLD | 99.9998 |
| Alkaline phosphatase (ALP) | Opt Width | 100 |
| Phosphate | Opt-HD | 87.1195 |
| Phosphate | Opt-LD | 85.4277 |

| Parameter | Deviation | Quantile 95% |
|---|---|---|
| Phosphate | Opt-VLD | 99.7274 |
| Phosphate | Opt Width | 99.9596 |
| Platelets | Opt-HD | 99.4953 |
| Platelets | Opt-LD | 98.0077 |
| Platelets | Opt-VLD | 99.9996 |
| Platelets | Opt Width | 100 |
| Basophils (abs) | Opt-HD | 100 |
| Basophils (abs) | Opt Width | 100 |
| Eosinophils (abs) | Opt-HD | 99.9999 |
| Eosinophils (abs) | Opt-LD | 99.9554 |
| Eosinophils (abs) | Opt-VLD | 99.9998 |
| Eosinophils (abs) | Opt Width | 100 |
| Neutrophils (abs) | Opt-HD | 91.9477 |
| Neutrophils (abs) | Opt-LD | 78.692 |
| Neutrophils (abs) | Opt-VLD | 96.0434 |
| Neutrophils (abs) | Opt Width | 99.9909 |
| Potassium | Opt-HD | 88.5189 |
| Potassium | Opt-LD | 86.9826 |
| Potassium | Opt-VLD | 99.8211 |
| Potassium | Opt Width | 99.8126 |
| Total protein | Opt-HD | 89.2193 |
| Total protein | Opt-LD | 90.9946 |
| Total protein | Opt-VLD | 99.9041 |
| Total protein | Opt Width | 99.9021 |
| Reticulocytes | Opt-HD | 99.8965 |
| Reticulocytes | Opt-LD | 98.2047 |
| Reticulocytes | Opt-VLD | 99.9972 |
| Reticulocytes | Opt Width | 100 |
| Sodium | Opt-HD | 97.2054 |
| Sodium | Opt-LD | 97.1906 |
| Sodium | Opt-VLD | 99.9999 |
| Sodium | Opt Width | 99.9953 |
| Testosterone | Opt-HD | 93.616 |
| Testosterone | Opt-LD | 90.0263 |
| Testosterone | Opt-VLD | 99.9997 |
| Testosterone | Opt Width | 100 |
| Thyroid-stimulating hormone (TSH) | Opt-HD | 99.8596 |
| Thyroid-stimulating hormone (TSH) | Opt-LD | 95.9359 |
| Thyroid-stimulating hormone (TSH) | Opt-VLD | 99.9729 |
| Thyroid-stimulating hormone (TSH) | Opt Width | 100 |
| Free thyroxine (FT4) | Opt-HD | 98.1266 |
| Free thyroxine (FT4) | Opt-LD | 96.016 |
| Free thyroxine (FT4) | Opt-VLD | 99.9937 |
| Free thyroxine (FT4) | Opt Width | 100 |
| Transferrin | Opt-HD | 99.7627 |
| Transferrin | Opt-LD | 99.4108 |
| Transferrin | Opt-VLD | 100 |
| Transferrin | Opt Width | 100 |
| Triglycerides | Opt-HD | 93.4211 |
| Triglycerides | Opt-LD | 74.3938 |
| Triglycerides | Opt-VLD | 93.289 |
| Triglycerides | Opt Width | 99.9987 |
| Free triiodothyronine (FT3) | Opt-HD | 95.0787 |
| Free triiodothyronine (FT3) | Opt-LD | 96.2396 |
| Free triiodothyronine (FT3) | Opt-VLD | 99.9999 |
| Free triiodothyronine (FT3) | Opt Width | 99.9999 |
| Uric acid | Opt-HD | 97.0569 |
| Uric acid | Opt-LD | 93.9664 |
| Uric acid | Opt-VLD | 99.9675 |
| Uric acid | Opt Width | 100 |
| Urea | Opt-HD | 91.3601 |
| Urea | Opt-LD | 83.0491 |
| Urea | Opt-VLD | 98.6482 |
| Urea | Opt Width | 99.9969 |
| Mean corpuscular volume (MCV) | Opt-HD | 99.9991 |
| Mean corpuscular volume (MCV) | Opt-LD | 99.9994 |
| Mean corpuscular volume (MCV) | Opt-VLD | 100 |
| Mean corpuscular volume (MCV) | Opt Width | 100 |
| Mean platelet volume (MPV) | Opt-HD | 99.9707 |
| Mean platelet volume (MPV) | Opt-LD | 99.8388 |
| Mean platelet volume (MPV) | Opt-VLD | 100 |
| Mean platelet volume (MPV) | Opt Width | 100 |

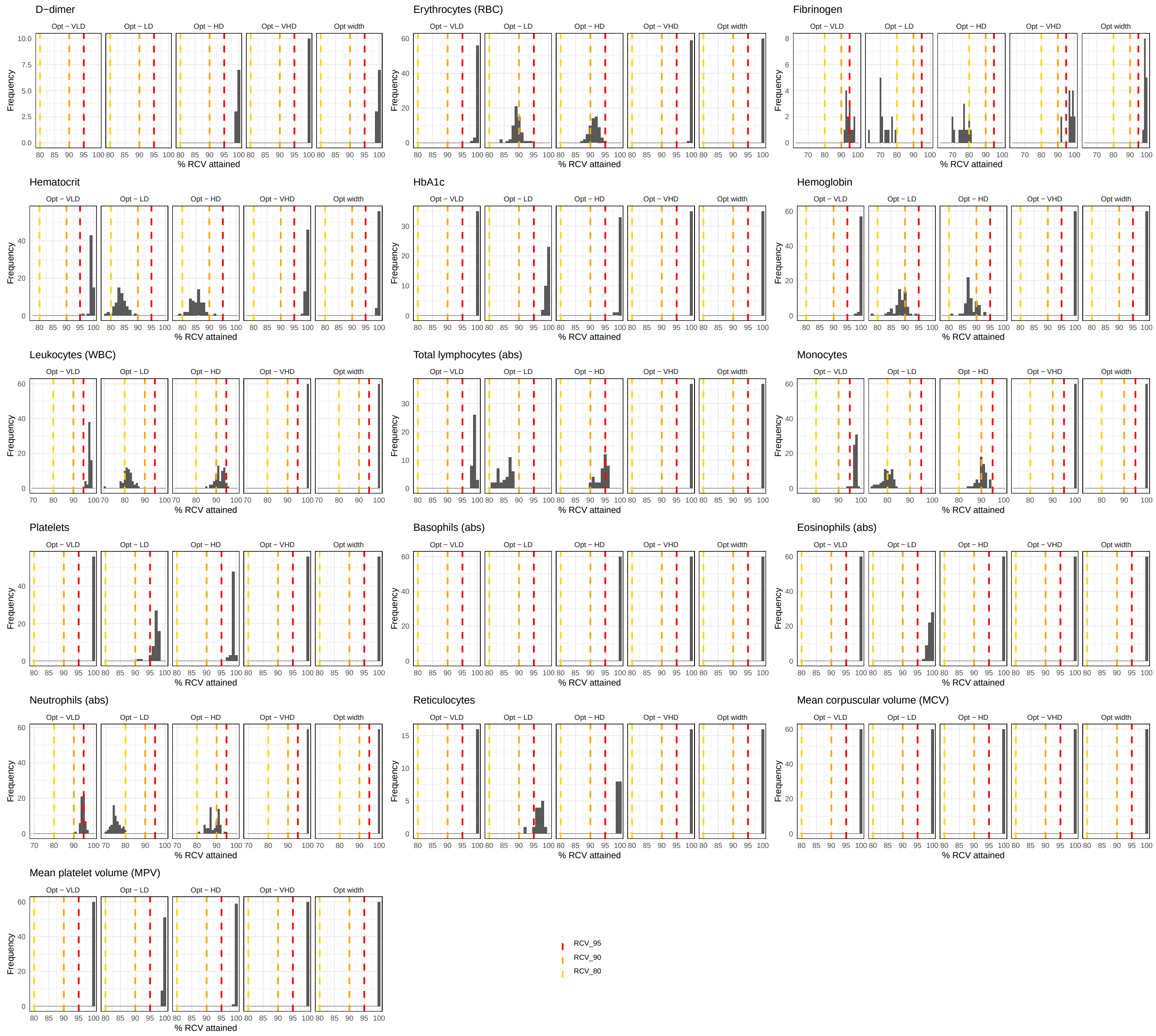


**Supplemental Fig. 1.** $p_{\mathrm{RCV}}$ for hematological biomarkers

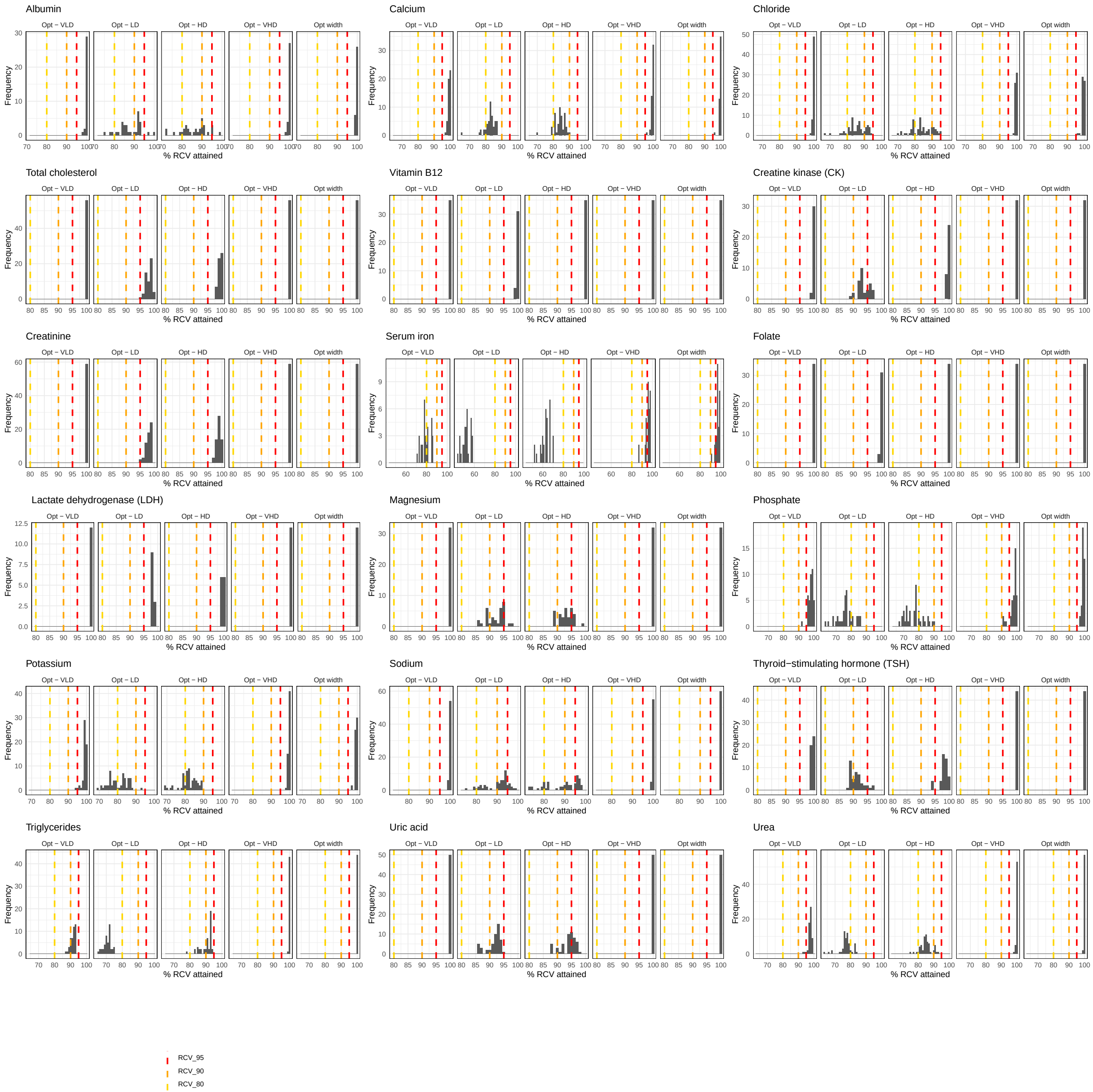


**Supplemental Fig. 2.** $p_{\mathrm{RCV}}$ for biochemistry biomarkers

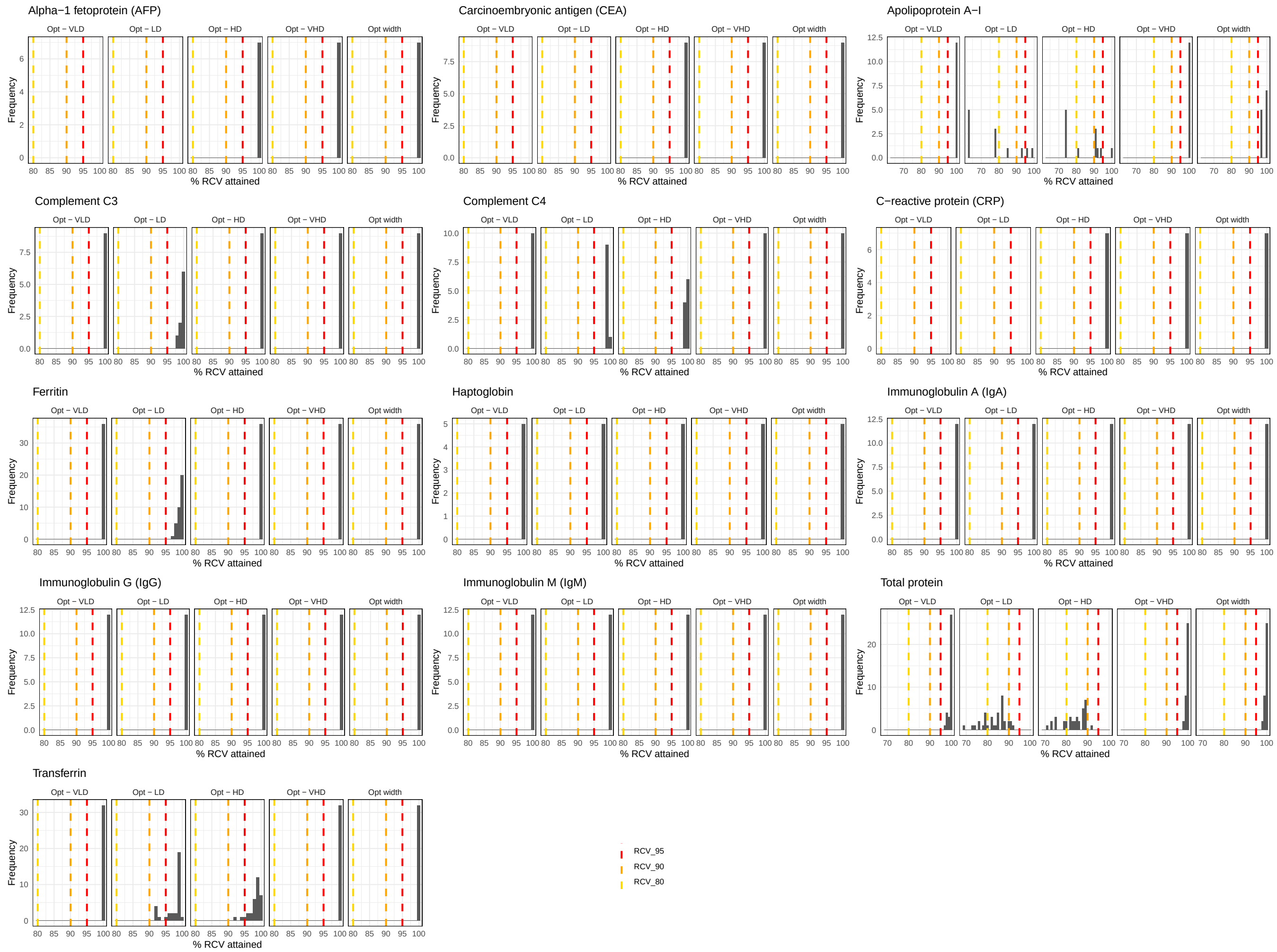


**Supplemental Fig. 3.** $p_{\mathrm{RCV}}$ for inflammatory, immulonological and cancer biomarkers

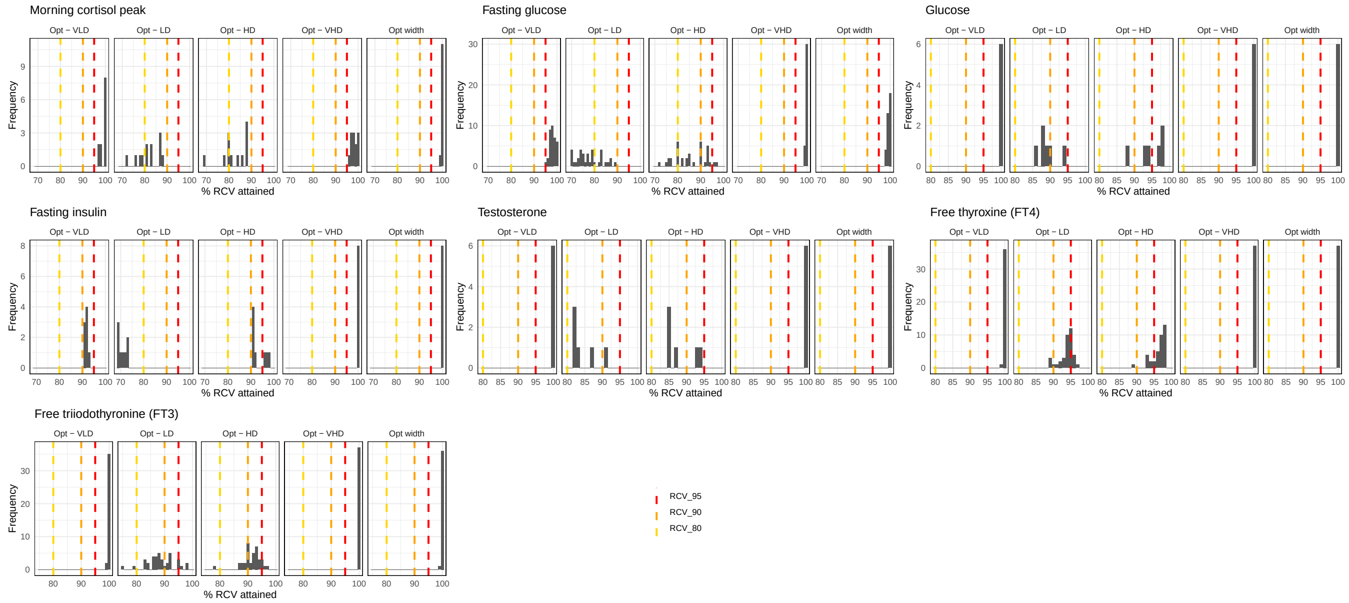


**Supplemental Fig. 4.** $p_{\mathrm{RCV}}$ for endocrinology and metabolism and biomarkers

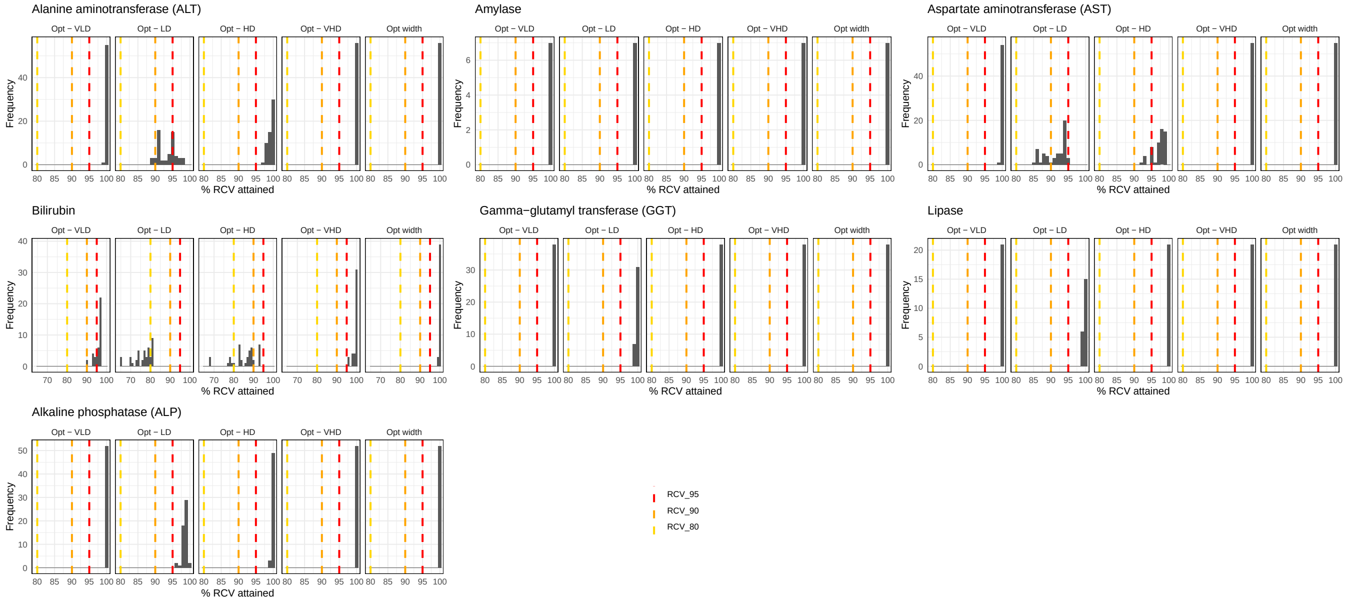


**Supplemental Fig. 5.** $p_{\mathrm{RCV}}$ for enzymes and liver biomarkers
